# Integrative Interactome and Ubiquitinome Analyses Reveal Multiple Regulatory Pathways Targeted by ICP0 in HSV-1 Infected Neuronal Cells

**DOI:** 10.1101/2021.12.13.472348

**Authors:** Fujun Hou, Zeyu Sun, Yue Deng, Siyu Chen, Xiyuan Yang, Feiyang Ji, Menghao Zhou, Keyi Ren, Dongli Pan

## Abstract

Herpes simplex virus 1 (HSV-1) is a neurotropic virus that can undergo both productive and latent infection in neurons. ICP0 is an HSV-1 E3 ubiquitin ligase crucial for productive infection and reactivation from latency. However, its targets have not been systematically investigated in neuronal cells. After confirming the importance of ICP0 in HSV-1 neuronal replication using an ICP0-null virus, we identified many ICP0-interacting proteins in infected neuronal and non-neuronal cells by mass-spectrometry-based interactome analysis. Co-immunoprecipitation assays validated ICP0 interactions with ACOT8, C1QBP, OTUD4, SNX9 and VIM in both Neuro-2a and 293T cells. Overexpression and knockdown experiments showed that SNX9 restricted replication of the ICP0-null but not wild-type virus in Neuro-2a cells. Ubiquitinome analysis by immunoprecipitating the trypsin digested ubiquitin reminant followed by mass spectrometry identified numerous candidate ubiquitination substrates of ICP0 in infected Neuro-2a cells, among which OTUD4 and VIM were novel substrates confirmed to be ubiquitinated by transfected ICP0 in Neuro-2a cells despite no evidence of their degradation by ICP0. Expression of OTUD4 was induced independently of ICP0 during HSV-1 infection. Overexpressed OTUD4 enhanced type I interferon expression during infection with the ICP0-null but not wild-type virus. In summary, by combining two proteomic approaches followed by confirmatory and functional experiments, we identified and validated multiple novel targets of ICP0 in neuronal cells, and revealed potential restrictive activities of SNX9 and OTUD4 as well as ICP0-dependent antagonism of these activities.

**Author Summary:** Herpes simplex virus 1 (HSV-1) establishes latent infection in neurons. ICP0 is known for its critical role in antagonizing cellular restrictive functions thereby initiating productive infection. It has been demonstrated to be important for both acute infection and reactivation from latency in neurons. However, little is known about its targets in neuronal cells. Here we combined two proteomic approaches, interactome and ubiquitinome analyses, to integratively identify interaction partners and substrates of ICP0 in HSV-1 infected neuronal cells. The results identified many novel targets as well as confirming previously reported ones. We also further validated some of the binding interactions and ubiquitin modifications. Functional studies revealed that the ICP0-interacting protein SNX9 restricted HSV-1 replication and the ICP0 substrate OTUD4 was induced to enhance type I interferon expression during HSV-1 neuronal infection. Moreover, the activities of these proteins appeared to be antagonized by ICP0-dependent mechanisms. This study provided comprehensive insight into ICP0 targets in neuronal cells and might prompt further investigation into the newly identified targets of ICP0.

## Introduction

Herpes simplex virus 1 (HSV-1) is a DNA virus that widely infects humans and can cause lesions in orolabial mucosa, eyes and genital areas during productive (lytic) infection. HSV-1 gene expression during lytic infection proceeds in a cascade fashion, with immediate-early, early and late genes being sequentially expressed. After lytic infection, HSV-1 establishes latent infection in sensory ganglia characterized by existence of viral genomes in neuronal nuclei with little protein expression but high expression of latency-associated transcripts. Latent virus can reactivate under certain stress to cause recurrent disease. This switch between lytic and latent infection enables the virus to permanently reside within the host while maintaining the ability to spread [1]. In rare incidences, the virus can travel from the peripheral nervous system to the central nervous system to cause life-threatening encephalitis. HSV-1 neuronal infection might also be related to chronic neurological disorders such as Alzheimer’s disease [2].

HSV-1 has evolved various strategies to evade the host’s defense mechanisms. One such strategy entails encoding infected cell protein 0 (ICP0), an immediate-early protein with E3 ubiquitin ligase activity. ICP0 functions via its really interesting new gene (RING) finger domain to ubiquitinate its substrates for proteosomal degradation. ICP0 is also a small ubiquitin-related modifier (SUMO)-targeted ubiquitin ligase, although not all its activity depends on SUMO modification [3,4]. ICP0 substrates have been reported to be involved in a wide range of host processes including intrinsic and innate immunity, chromatin remodeling and DNA damage response (DDR) pathways [5–8] (reviewed in [9–11]). For example, ICP0-induced degradation of PML leads to the disruption of PML nuclear bodies and release of entrapped viral genomes [12–18]. ICP0 induces the degradation of chaperone proteins of histone H3.3 including IFI16 and ATRX resulting in attenuation of epigenetic silencing of viral genes [19–22]. ICP0 can also bind to CoREST and block CoREST-mediated silencing of viral genes by dissociating HDAC1 [23]. ICP0 targeting of RNF8 and RNF168, which are ubiquitin ligases mediating H2A and H2AX ubiquitination in the DDR pathway, leads to failure to recruit host DDR repair factors [6,24]. ICP0 also scrambles innate immune pathways through degradation of DNA-PKcs [25]. Because of these mechanisms that antagonize host repressive functions, ICP0 is important for efficient initiation of lytic infection, and replication of HSV-1 is severely impaired without ICP0 or its RING finger domain in multiple cell types especially at low multiplicities of infection (MOIs) [26,27].

High-throughput proteomic approaches have been successfully employed to discover host interacting proteins and substrates of ICP0. One study characterized changes in cellular SUMO2 proteome upon HSV-1 infection in hepatocytes [28]. A recently published study compared wild-type virus and a mutant lacking functional ICP0 in the proteome associated with viral DNA [29]. We also notice three studies of the ICP0 interactome, one in ICP0-transfected 293T cells [30], one in HSV-1 infected Hep-2 cells [31] and one in HSV-1 infected 293T cells [32]. However, we are unaware of any proteomic research specifically on ICP0 in neuronal cells.

Being the site of HSV latency and reactivation, the nervous system is highly relevant to the HSV life cycle. There is much evidence that ICP0 is important for HSV-1 neuronal infection. For example, ICP0 mutant viruses were severely impaired in establishment of latency and reactivation in ganglionic neurons or tissues [33–35] and delivery of ICP0 by adenovirus to ganglionic neurons stimulated HSV-1 reactivation from latency [36]. An HSV-1 mutant with increased ICP0 expression due to mutations in binding sites of a neuron-specific miRNA showed increased overall lytic gene expression in acutely and latently infected mouse trigeminal ganglia [37,38]. ICP0 deficiency also affected the levels of heterochromatin on viral genes in latently infected ganglia [39]. However our knowledge about the mechanisms by which ICP0 exerts these important functions was quite limited. Therefore we undertook this study to systematically investigate into ICP0 targets in neuronal cells. Besides the commonly performed interactome analysis, comprehensive protein ubiquitination investigation for this ubiquitin E3 ligase should greatly facilitate target identification. Because ubiquitin forms an isopeptide bond with its substrate and generates a signature di-glycine modification on lysine following trypsin digestion, immunoprecipitation using di-glycine antibody has been widely applied to enrich peptides with ubiquitin remnants before mass spectrometry (MS)-based shot-gun proteomics survey of ubiquitinated proteins (hereafter referred to as ubiquitinome) [40,41]. Recently, di-glycine enrichment has been coupled with tandem mass tag (TMT)-based multiplexed isotopic labeling for parallel quantitative ubiquitinome analysis [42,43]. In this work, we employed this method in combination with immunoprecipitation-based interactome analysis to identify ICP0 targets in neuronal cells followed by assessing the effects of these targets on viral replication and the host interferon response.

## Results

### ICP0 was confirmed to be important for HSV-1 replication in primary neurons and Neuro-2a cells

To better understand the importance of ICP0 in HSV-1 neuronal replication, we first examined replication of an ICP0-null virus, 7134, relative to its rescued derivative, 7134R in cultured primary neurons isolated from mouse trigeminal ganglia (TG). The isolation was performed with density gradient centrifugation following dissociation of the tissues to obtain relatively pure neurons [44]. Viral titers were determined by plaque assays in U2OS cells, which are permissive for ICP0-null virus replication. After infection of the primary neurons, the supernatant titers of 7134 virus were ~2 logs lower than those of 7134R virus at both MOIs of 1 and 10 from 24 to 72 h post-infection (hpi) (Fig. 1A). For proteomic analysis, which requires a large number of cells, we considered using Neuro-2a cells derived from mouse brain neuroblastoma cells commonly used in many HSV-related studies [37,45,46]. After infection of Neuro-2a cells, 7134 virus replicated to ~2 logs lower titers than 7134R at MOIs of 0.1 and 1 at 18 hpi (Fig. 1B). However the defect of 7134 virus was modest at the high MOI of 10 in agreement with the MOI dependence of ICP0 effects commonly observed in cell lines [26,27]. Under the same conditions, we also compared 7134 and 7134R in HFF (human foreskin fibroblast) and 293T (human embryonic kidney) cells. The defects of 7134 virus were more severe in HFF cells but moderate in 293T cells consistent with the known cell-type specific role of ICP0. These results substantiated the importance of ICP0 in HSV-1 neuronal replication and suggested that Neuro-2a cells could be used to investigate the important interactions with ICP0 in neuronal cells.

**Fig. 1.**
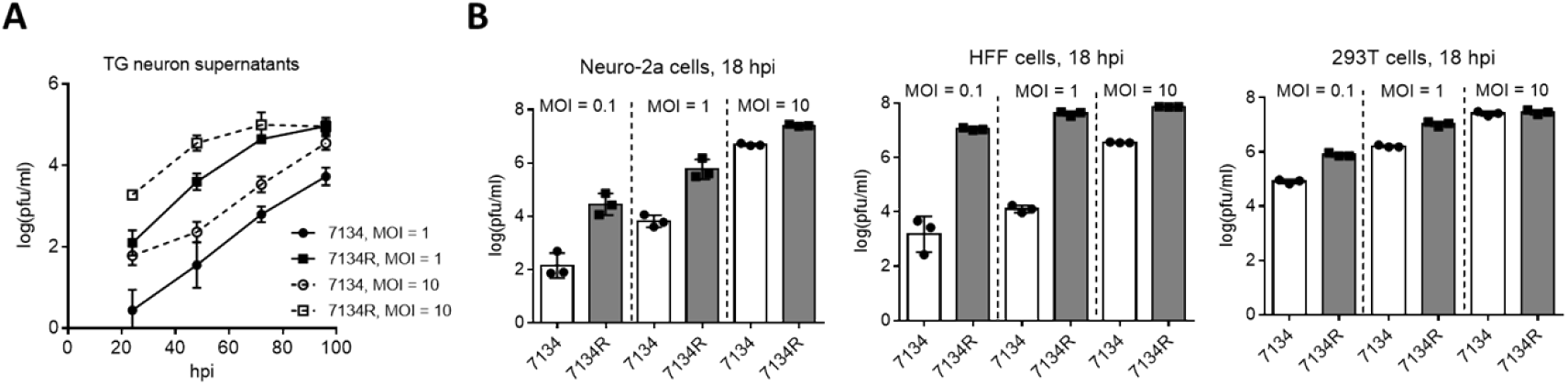
ICP0 is important for HSV-1 replication in neuronal cells. (A) Primary mouse TG neurons were infected with 7134 or 7134R at an MOI of 1 or 10 as indicated. At the indicated times post-infection, supernatants were collected for virus titration. (B) Neuro-2a (left), HFF (middle) and 293T (right) cells were infected with 7134 or 7134R at the indicated MOI and harvested at 18 hpi for virus titration. Data are presented as mean values ± standard deviations (S.D.).

### Construction and characterization of Flag-ICP0 virus

To facilitate interactome analysis in infected cells, we constructed Flag-ICP0 virus that expresses N-terminal Flag-tagged ICP0 using bacterial artificial chromosome (BAC) technology on the basis of wild-type (WT) BAC virus described previously (here referred to as WT virus) (Fig. 2A) [47]. The Flag-tag expressing sequence was inserted into both copies of the ICP0 gene. The resulting Flag-ICP0 virus replicated with WT kinetics in Vero and Neuro-2a cells (Fig. 2B). A Flag antibody readily detected ICP0 expression from the virus in 293T, HFF and Neuro-2a cells (Fig. 2C). ICP0 expression reached plateaus in all the three cell lines at around 6 hpi, so we decided to use 6 h as the time point to collect samples for the following interactomic experiments.

**Fig. 2.**
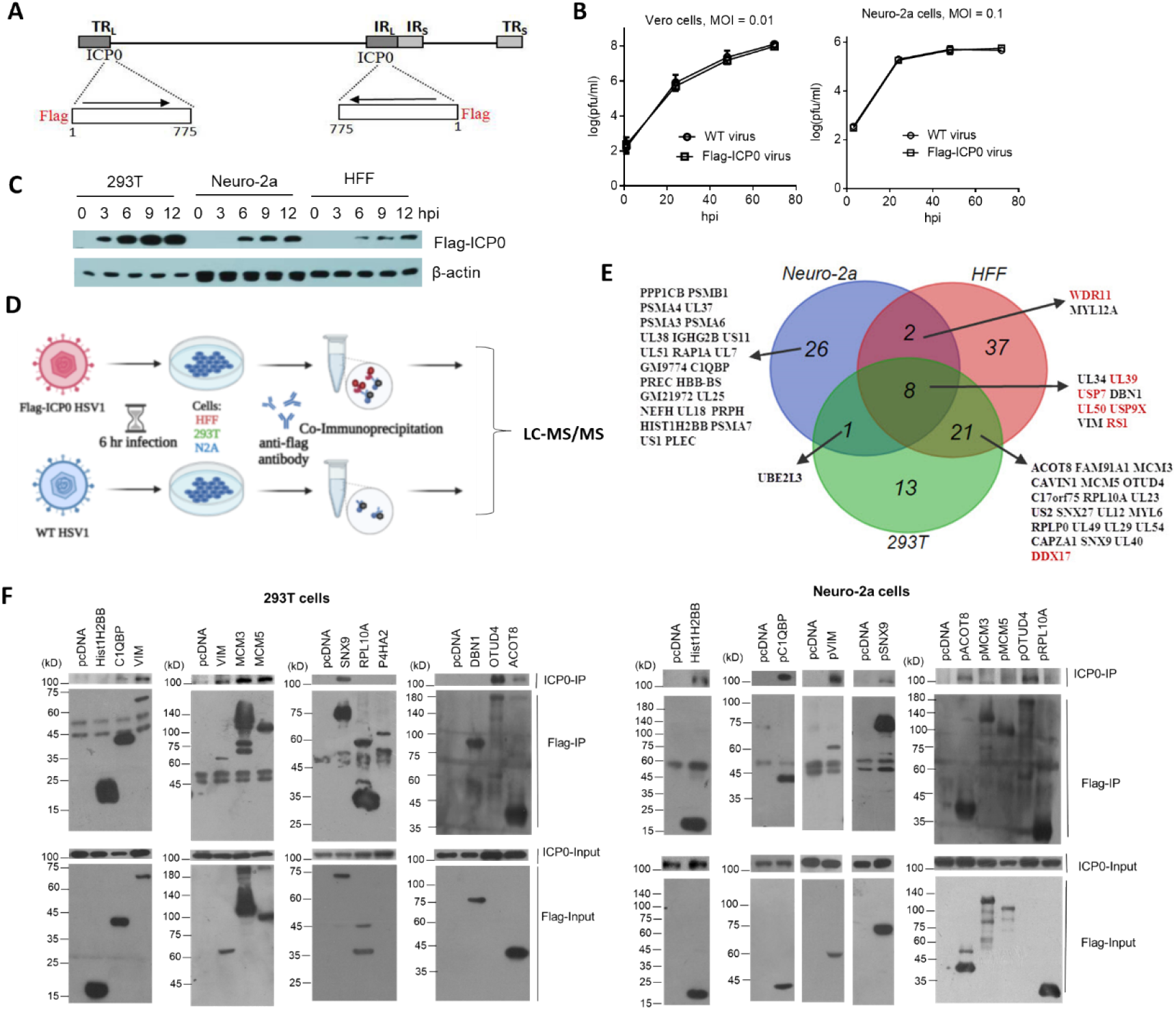
ICP0-interactome analysis. (A) Diagram of recombinant Flag-ICP0 virus. The horizontal line at the top represents HSV-1 genome and the boxes labeled with TR_L_, IR_L_, TR_S_ and IR_S_ represent the repeated sequences. At the bottom are expanded ICP0 gene regions with the inserted N-terminal Flag tags indicated. (B) Virus growth curves of WT and Flag-ICP0 virus in Vero (left) and Neuro-2a (right) cells after infection at an MOI of 0.01 and 0.1, respectively. Data are presented as mean values ± S.D. (C) Neuro-2a, HFF and 293T cells were infected with Flag-ICP0 virus at an MOI of 10 and cells were lysed at the indicated time points for examination of Flag-ICP0 expression by western blots with anti-Flag and anti-β-actin antibodies. (D) Schematic diagram of the procedures for ICP0 interactome analysis. Neuro-2a, 293T and HFF cells were infected with WT (control) or Flag-ICP0 virus for 6 h at an MOI of 20 with three biological replicates. Proteins were immunoprecipitated by the anti-Flag antibody and examined by LC-MS/MS. (E) Venn diagram showing candidate ICP0-interacting proteins identified by the interactome analysis in the three cell types. Previously reported ICP0-interacting proteins are marked in red. (F) 293T (left) or Neuro-2a (right) cells in each 100-mm plate were co-transfected with 7.5 μg of an empty vector (pcDNA) or a plasmid expressing the indicated protein linked to a Flag-tag and 7.5 μg of a plasmid expressing untagged ICP0 for 48 h. After the cells were lysed, the indicated proteins were immunoprecipitated by an anti-Flag antibody and analyzed by western blots for ICP0 and the corresponding proteins using an ICP0 and Flag antibody, respectively. Results from immunoprecipitated samples (IP) were shown in the upper panels and those from the lysates (Input) were shown in the lower panels.

### ICP0 interactome analysis in HSV-1 infected cells

To analyze the ICP0 interactome, we infected Neuro-2a, HFF and 293T cells with Flag-ICP0 or WT virus for 6 h before co-immunoprecipitation (Co-IP) with a Flag antibody followed by protein identification by liquid chromatography-mass spectrometry/mass spectrometry (LC-MS/MS) (Fig. 2D). HFF and 293T cells were used to learn about the cell-type specificity of the interactome. After applying selection criteria (Materials and Methods), we identified 37, 68 and 43 potential ICP0-interacting proteins in Neuro-2a, HFF and 293T cells, respectively, amounting to a total of 108 candidate proteins (Fig. 2E, Table S1). Functional enrichment analysis revealed that ICP0 interacts with a host protein network related to several biological function clusters including deubiquitination, proteasome, cytoskeleton, translation elongation, mRNA processing and protein transport (Fig. S1). Many previously reported ICP0 targets were also confirmed in these data, including viral proteins UL50 [32], UL39 [32] and RS1 (ICP4) [48,49] as well as host proteins USP7 [50], USP9X [31], PRKDC [51], DDX17 [52] and WDR11 [8]. Interestingly, eight proteins were found in all three cell lines: viral proteins UL50, UL39, RS1 and UL34, and host proteins USP7, USP9X, VIM and DBN1. Twenty-six proteins were identified exclusively in Neuro-2a cells (Fig. 2E) suggesting that they might represent neuron-specific interactions.

### Validation of novel interactions with ICP0

To validate the proteomics data, we selected some top novel host candidates according to Flag-ICP0/WT ratios. Priorities were given to the candidates identified in Neuro-2a cells. However, considering that results from the mouse cells might miss interactions occurring in human cells, we also included some top candidates identified in both 293T and HFF cells. Thus, we selected a total of 17 top candidates and cloned their human genes into expressing plasmids with Flag tags. After co-transfection of plasmids with an ICP0 expressing plasmid into Neuro-2a and 293T cells, we performed IP in a reverse way relative to the interactome analysis. We confirmed that untagged ICP0 could be pulled down by Flag-tagged C1QBP, VIM, SNX9, ACOT8 and OTUD4 in both Neuro-2a and 293T cells (Fig. 2F). We note that although SNX9, ACOT8 and OTUD4 were not identified in the interactome analysis in Neuro-2a cells possibly due to limited sensitivity of mass spectrometry or the human-mouse differences, the reverse Co-IP showed that their human forms were all able to interact with ICP0 in Neuro-2a cells. Therefore, these proteins were included in the following functional studies. Hist1H2BB co-precipitated with ICP0 in Neuro-2a but not 293T cells. MCM3 and MCM5 co-precipitated with ICP0 in 293T but not Neuro-2a cells. Other proteins that we examined, including HNRNPM, P4HA2, RPL10A, MYL12A, PPP1CB, CAVIN1, DBN1, PSMA3 and PSMA6, could not reproducibly co-immunoprecipitate with ICP0 in either cell line (data not shown).

### ICP0-interacting protein SNX9 restricted replication of an ICP0-null virus in Neuro-2a cells

To explore the functions of the ICP0-interacting proteins, we focused on the five proteins that demonstrated robust interactions with ICP0 in both 293T and Neuro-2a cells, including C1QBP, VIM, SNX9, ACOT8 and OTUD4. Besides using the plasmids to overexpress the proteins, we designed at least two siRNAs to knock down each protein. After evaluating the knockdown efficiencies, two siRNAs with the highest efficiencies for each protein was selected along with two negative control siRNAs. Efficient knockdown by the selected siRNAs and overexpression by the plasmids in Neuro-2a cells were confirmed in western blots (Fig. S2). We then infected the cells after transfection of the plamids or siRNAs. Overexpression of none of these proteins had a significant effect on replication of 7134 or KOS virus (Fig. 3A). However both siRNAs against SNX9 resulted in significantly increased 7134 virus titers relative to both control siRNAs in Neuro-2a cells, although only one of them (SNX9-si1) showed modestly increased KOS titers (Fig. 3B). We then performed co-transfection of the siRNAs together with plasmids. While both siRNAs against SNX9 significantly increased 7134 virus titers in the presence of the control plasmid, they could not do so in the presence of the SNX9 expressing plasmid (Fig. 3C) suggesting that the increased replication was caused by specific knockdown of SNX9. Growth curve analysis showed that even the less effective SNX9 siRNA (SNX9-si2) significantly increased replication kinetics of 7134 virus at an MOI of 0.2 but it had no effect at an MOI of 5 (Fig. 3D). Therefore endogenous SNX9 restricted HSV-1 replication at a low MOI.

**Fig. 3.**
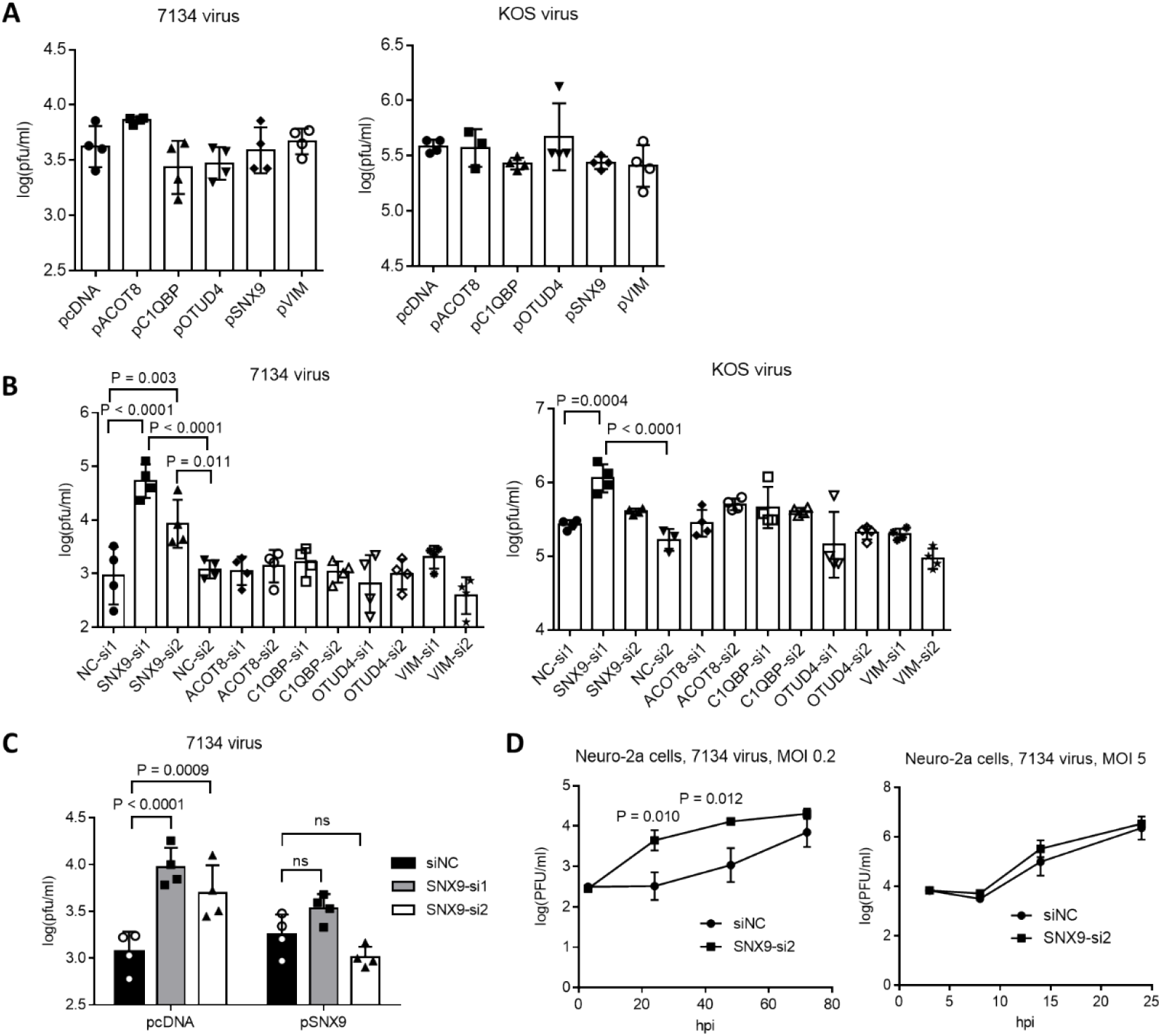
Effects of ICP0-interacting proteins on HSV-1 replication in Neuro-2a cells. (A) Neuro-2a cells were transfected with 400 ng/ml of the indicated plasmids for 40 h, then infected with 7134 (left) or KOS (right) virus for 48 h at an MOI of 0.2 before the cells were harvested for virus titration by plaque assays. (B) Neuro-2a cells were transfected with 80 nM of the indicated siRNAs for 40 h, then infected with 7134 (left) or KOS (right) virus for 48 h at an MOI of 0.2 before the cells were harvested for virus titration. NC, negative control. (C) Neuro-2a cells were transfected with 80 nM of the indicated siRNAs for 17 h and then transfected with 600 ng/ml of the indicated plasmids for 33 h before infection with 7134 virus at an MOI of 0.2 for 44 h. The cells were then harvested for viral titration. (D) Neuro-2a cells were transfected with a negative control siRNA or an siRNA against SNX9 for 40 h before infection with 7134 virus at an MOI of 0.2 (left) or 5 (right). The cells were harvested at the indicated times for virus titration. Data were analyzed by one-way ANOVA with Donnett’s multiple comparisons tests (A, B), two-way ANOVA with Sidak’s multiple comparisons tests (C), or two-tailed unpaired t-tests (D). Data are presented as mean values ± S.D.

### ICP0 ubiquitinome analysis in infected Neuro-2a cells

To identify ubiquitination substrates of ICP0, we compared the ubiquitinome during HSV-1 infection in the presence versus absence of ICP0. Absence of ICP0 was achieved using the ICP0-null virus 7134, which was compared with 7134R. To increase the stringency of the experiment, we also considered rescuing the absence of ICP0 by transfection. Therefore we designed the following three groups: group A, control transfection + 7134 virus infection; group B, ICP0 transfection + 7134 virus infection; group C, control transfection + 7134R virus infection. Only ubiquitination events that were induced both by transfected ICP0 (comparing B and A) and by ICP0 expressed from the virus (comparing C and A) were considered, which should greatly reduce frequencies of false-positive discoveries. Neuro-2a cells were transfected for 24 h and then infected for 6 h at an MOI of 20 before total protein was digested by trypsin and immunoprecipitated with ubiquitin remnant K-ε-GG antibody followed by TMT labeling and LC-MS/MS analysis (Fig. 4A). Overall, ratios of protein quantities in group B versus A (B/A) and group C versus A (C/A) comparisons correlated well with each other (Fig. 4B). Ubiquitinated sites potentially modified by ICP0 were selected by applying criteria of either false discover rate (FDR) < 0.05 or fold change > 1.25 in both comparisons. After mapping these sites to proteins, we identified 1022 proteins in the B/A comparison and 522 proteins in the C/A comparison. The 351 proteins identified by both B/A and C/A comparisons were considered as potential ICP0 substrates (Fig. 4C and Table S1). Functional enrichment analysis showed that the potential host substrates are enriched in multiple pathways, including metabolism of RNA/proteins, rRNA processing, cell cycle and deubiquitination etc. (Fig. 4C). Nine proteins were previously reported to be potential host substrates of ICP0 including USP7 [53], PML [54], OPTN [55], SUMO2 [28,56], UBE2E1 [57], ZBTB10 [28], MORC3 [58], RNF168 [6] and ETV6 [28]. Their potential ubiquitination sites determined by our mass spectrometry data are listed in Fig 4C. Additionally, the candidate substrate proteins were clustered in several superfamilies classified by conserved domains such as nucleoside triphosphate hydrolase, RNA helicase, Zinc-finger, RNA binding and ubiquitin (Table S3). Regarding viral proteins, out of 55 viral proteins with detected ubiquitination, 7 proteins, UL12, UL21, UL5, UL51, UL54, UL9 and US3 showed increased ubiquitination in the presence of ICP0 (Fig. S3), indicating that they might be potential viral substrates of ICP0.

**Fig. 4.**
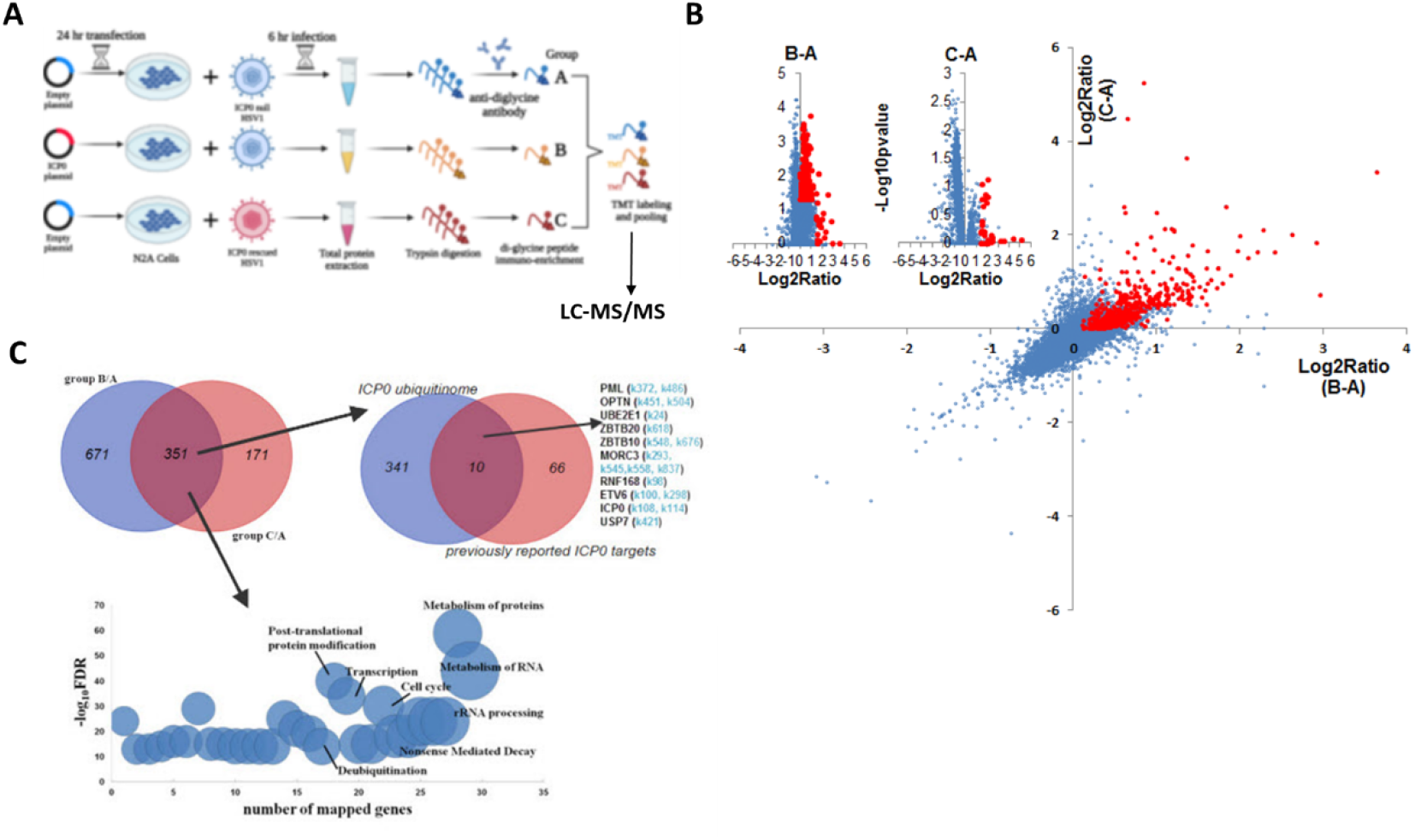
ICP0 ubiquitinome analysis in Neuro-2a cells. (A) Schematic diagram of the experimental design for ICP0 ubiquitinome analysis. (B) Potential ICP0 targeted sites (red) showing higher ubiquitination levels in group B than A (x-axis) or those showing higher ubiquitination levels in group C than A (y-axis) according to quantitative mass spectrometry. Individual volcano diagrams showing B-A and C-A comparisons were also displayed in the upper left corner. (C) Potential ICP0 ubiquitination target proteins identified by comparing groups B and A as well as comparing groups C and A. The overlap of the two comparisons represents 351 candidate targets for subsequent analysis. The pathways enriched in these proteins are shown in the bottom graph. The upper right Venn diagram shows the overlap of our ICP0 ubiquitinome results and previously reported ICP0 substrates or interacting proteins. For the targets in our data that were also previously reported, potential ubiquitination sites on lysine differentially represented between ICP0-positive and negative cells in our data are indicated in light blue.

### OTUD4 and VIM were ubiquitinated by ICP0 in Neuro-2a cells

Comparison of the interactome and ubiquitinome data identified 3 viral and 8 host proteins which both interacted with ICP0 and exhibited elevated ubiquitination in ICP0’s presence indicating that they were highly likely to be substrates of ICP0 (Fig. 5A). All of these are novel potential substrates of ICP0 except for the deubiquitinase USP7, whose ubiquitination by ICP0 has been reported elsewhere [53,59]. Because host proteins VIM and OTUD4 were confirmed to robustly interact with ICP0 (Fig. 2F), we focused on these two proteins for ubiquitination analysis. VIM was previously documented to be decorated with 11 SUMO2 modifications by another large-scale MS study [60]. Our MS data detected 17 potential ubiquitination sites in VIM (Fig. 4B). We note that the trypsin digestion resulted in different reminants in ubiquitinated or SUMOylated lysine, so the current LC-MSMS analysis was specific to ubiquitin modification. Interestingly the majority of the newly identified ubiquitination sites overlapped with the SUMOylation sites, indicating possible crosstalk between ubiquitination and SUMOylation in VIM. Almost all ubiquitination and SUMOylation sites are located in the functional coil regions of VIM. Notably, the only site (K282) showing increased ubiquitination due to ICP0 presence has not been reported to be SUMOylated. The MS data also detected 4 potential ubiquitination sites in OTUD4, all of which are near the OTU domain and not far from the previously annotated phosphorylation sites. One of the sites (K264) showed increased ubiquitination due to ICP0 presence and therefore could be an ICP0 target site. To examine whether VIM and OTUD4 were ubiquitinated by ICP0, we co-transfected ubiquitin, VIM or OTUD4 and ICP0 or its control (empty vector or ring finger mutant) into Neuro-2a cells before immunoprecipitation of VIM and OTUD4 and analysis of ubiquitination levels by western blots. Both VIM and OTUD4 exhibited substantial enhancement of ubiqutination in the presence of ICP0 relative to the respective controls suggesting that they could be ubiquitinated by ICP0 (Fig. 5C). However transfected ICP0 did not cause decreases in VIM and OTUD4 levels indicating that ubiquitination of these proteins may not lead to degradation.

**Fig. 5.**
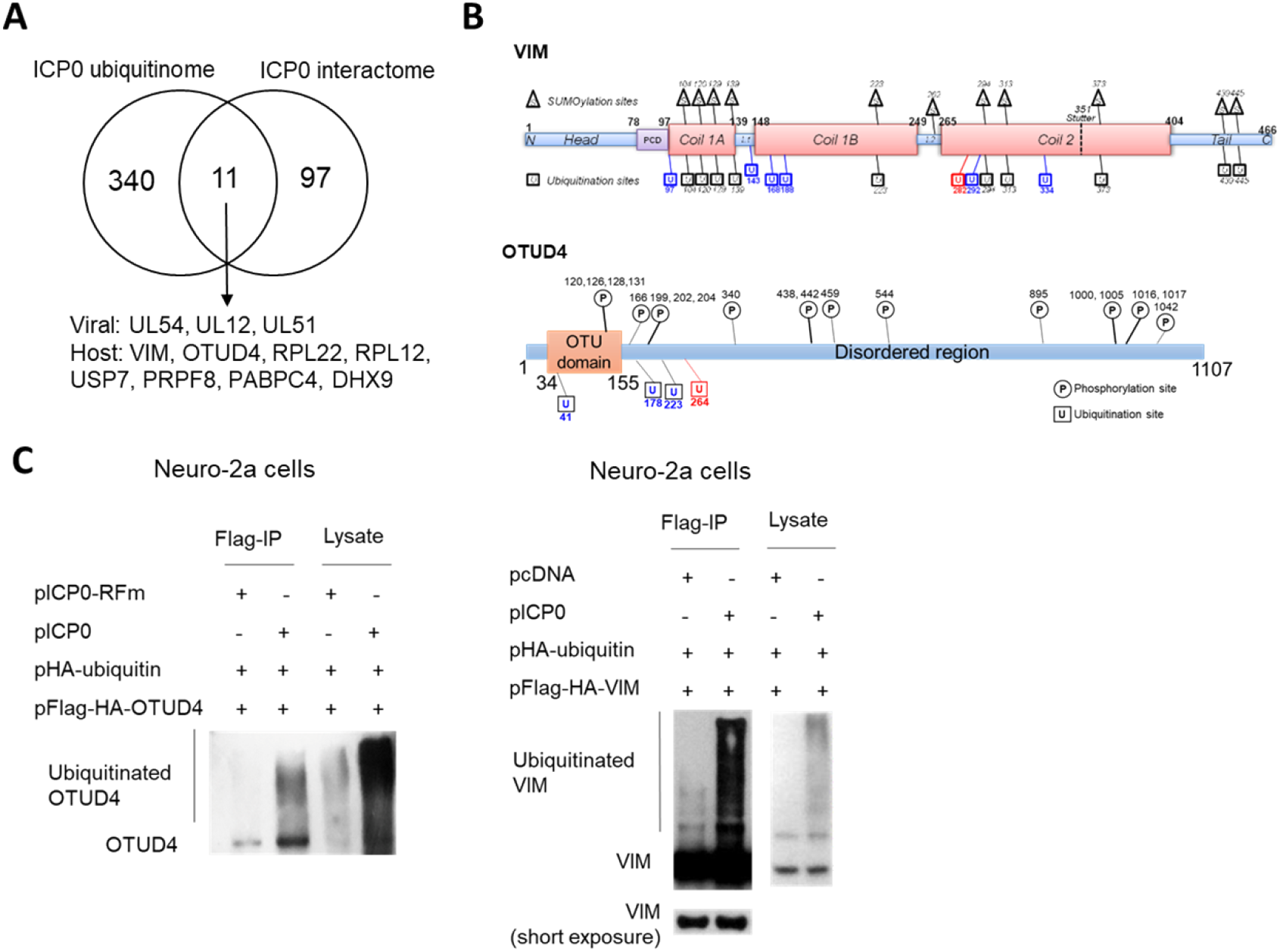
ICP0-mediated ubiquitination of OTUD4 and VIM. (A) Venn diagram showing 12 proteins identified by both ICP0 interactome and ubiquitinome analyses. (B) Summary of ubiquitination sites on VIM (upper) and OTUD4 (lower). Schematic representation of functional domains of VIM and OTUD4 decorated with previously documented SUMOylation (for VIM, in triangles) or phosphorylation (for OTUD4, in circles) sites and newly identified ubiquitination sites (in squares). L: linker region; PCD: pre-coiled domain. Blue indicates sites with ubiquitination but not SUMOylation. Red indicates the ubiquitination site potentially targeted by ICP0. (C) Neuro-2a cells in each 100-mm plate were co-transfected with 5 μg of a plasmid expressing Flag-HA-tagged OTUD4 (left) or VIM (right), 2 μg of a plasmid expressing HA-tagged ubiquitin, and 3 μg of a plasmid expressing ICP0 or its RING finger mutant (RFm) or an empty vector (pcDNA) for 48 h. Cell lysates were immunoprecipitated by an anti-Flag antibody. The lysates and immunoprecipitated samples (IP) were analyzed by western blots for ubiquitination levels by an anti-HA antibody.

### OTUD4 was upregulated to enhance type I interferon (IFN) expression during ICP0-null virus infection

To understand the roles of VIM and OTUD4 during HSV-1 infection, we first assessed their levels in infected Neuro-2a cells. Interestingly, we observed substantial upregulation of OTUD4, but not VIM starting from as early as 3 hpi (Fig. 6A). This upregulation was independent of the E3 ligase activity of ICP0 since it was also observed during infection with an ICP0 ring finger mutant (RFm) virus. The increases in OTUD4 protein levels correlated with similar increases in its mRNA levels indicating stimulation of *Otud4* gene transcription during infection (Fig. 6B). Comparison of WT and RFm viruses showed no indication of ICP0-induced degradation of OTUD4 or VIM at later times (Fig. 6A). Thus, OTUD4 was upregulated during HSV-1 infection without being degraded by ICP0.

**Fig. 6.**
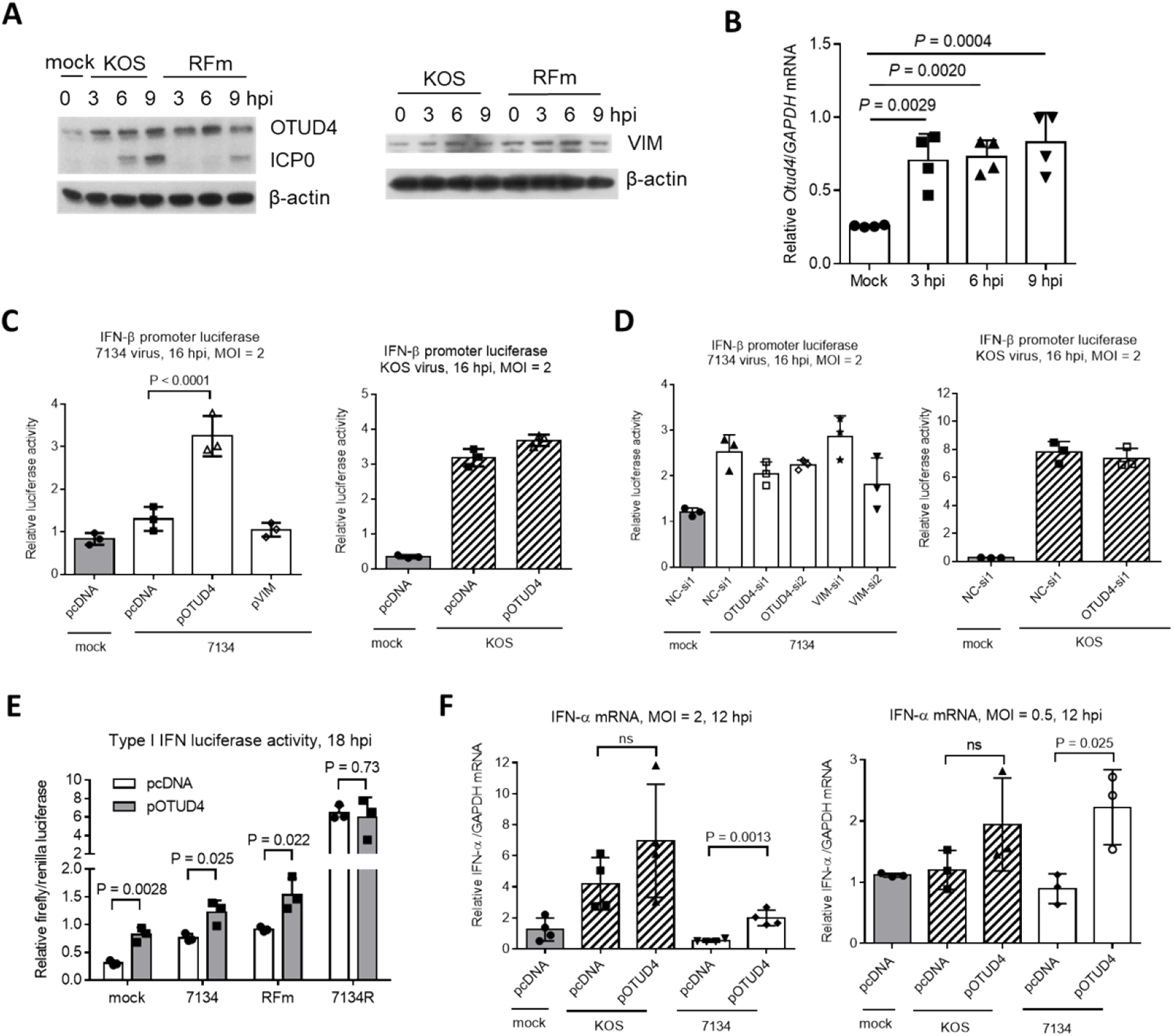
Functions of OTUD4 and VIM in Neuro-2a cells. (A) Cells were infected with KOS or ICP0 RING finger mutant virus (RFm) at an MOI of 20 for the indicated times before the cells were harvested for western blot analysis for the indicated proteins. (B) After infection of cells with KOS at an MOI of 20, the cells were harvested at the indicated times for qRT-PCR analysis of Otud4 mRNA levels normalized to GAPDH mRNA levels. (C) Cells were co-transfected with 400 ng/ml of the indicated plasmid, 80 ng/ml of a pIFN-β (an IFN-β promoter luciferase plasmid) and 40 ng/ml of pRL-TK (a control plasmid expressing renilla luciferase) for 48 h, then infected with 7134 (left) or KOS (right) at an MOI of 2 for 16 h before measurement of luciferase activities. (D) Neuro-2a cells were co-transfected with 80 nM of the indicated siRNA, 80 ng/ml of a pIFN-β and 40 ng/ml of pRL-TK for 48 h, then infected with 7134 (left) or KOS (right) at an MOI of 2 for 16 h before measurement of luciferase activities. (E) Same as C except for the different viruses used. (F) Cells were transfected with 400 ng/ml of the indicated plasmid, then infected with the indicated virus at an MOI of 2 (left) or 0.5 (right) for 12 h before qRT-PCR quantification of *IFN-α* mRNA normalized to *GAPDH* mRNA. Data were analyzed by one-way ANOVA with Donnett’s multiple comparisons tests (B, C, D, F) or two-tailed unpaired t-tests (E). Data are presented as mean values ± S.D.

To learn more about the roles OTUD4 and VIM, although they showed no effect on HSV-1 replication (Fig. 3A, B), we wondered whether they could regulate innate immune response to infection. To test this, we first transfected Neuro-2a cells with an IFN-β promoter luciferase reporter construct together with a plasmid expressing OTUD4 or VIM before infection at an MOI of 2 for 16 h. 7134 virus only modestly stimulated luciferase expression whereas KOS virus showed much stronger stimulation. In 7134 infected cells, overexpression of OTUD4 significantly increased luciferase activity whereas knocking down OTUD4 or overexpressing or knocking down VIM had no significant effect (Fig. 6C, D). However, in KOS infected, neither overexpressing nor knocking down OTUD4 had a significant effect indicating an ICP0-dependent block of OTUD4 activity. Consistent with this, overexpressed OTUD4 increased luciferase activity during infection with an ICP0 RFm virus but had no effect during infection with 7134R virus (Fig. 6E). Unexpectedly, OTUD4 also increased luciferase activity during mock infection suggesting that OTUD4 could raise the baseline expression of IFN-β in unstimulated cells. In concordance with the luciferase results, quantification of endogenous IFN-α mRNA showed that induction of IFN-α expression by KOS but not 7134 virus was obvious at an MOI of 2, and overexpressed OTUD4 significantly increased IFN-α mRNA levels in 7134 but not KOS infected cells (Fig. 6F). At a lower MOI of 0.5, where induction of IFN-α expression was no longer detectable even in KOS infected cells, overexpressed OTUD4 significantly increased IFN-α mRNA levels in 7134 but not KOS virus infected cells. Therefore OTUD4 could enhance type I IFN expression during HSV-1 infection and this function appeared to be blocked by ICP0-dependent processes.

## Discussion

After confirming that ICP0 is important for HSV-1 lytic infection in neuronal cells, we combined quantitative interactome and ubiquitinome approaches to identify targets of ICP0 in neuronal cells followed by validation of the interaction and ubiquitination events. Functional studies on the validated targets established SNX9 as a restriction factor and OTUD4 as a regulator of type I IFN production and provided evidence that the functions of these proteins might be modulated in an ICP0-dependent manner.

The use of two proteomic approaches allowed us to dissect binding and ubiquitination events, which may not always be coupled. Binding interactions may function in ways independent of ubiquitination, and some ubiquitin modifications might result from transient interactions that were not detected after immunoprecipitation. The ubiquitination analysis was also useful in identifying ubiquitination sites. Moreover, combining the two approaches should facilitate identification of valid substrates that were unlikely to be false-positives. Indeed we confirmed that both OTUD4 and VIM could be ubiquitinated by ICP0.

Our data recapitulated many of the previous reported ICP0-interacting proteins or substrates. Two important known substrates are USP7 and PML. While USP7 was identified in both interactome and ubiquitinome analyses, PML was identified only by the ubiquitinome analysis indicating that the interaction between ICP0 and PML might be too transient to be caught by immunoprecipitation. USP7 is a deubiquitinase that can inhibit ICP0 autoubiquitination thereby leading to ICP0 stabilization [3,53]. The role of PML in restriction of viral gene transcription and its antagonism by ICP0 have been well documented [12,14,18]. Our data suggest that these processes likely occur in neuronal cells too.

Functional studies on selective novel interaction partiners and/or substrates of ICP0 revealed two potential host restrictive factors as SNX9 restricted HSV-1 replication and OTUD4 could induce type I IFN production which might potentially restrict HSV-1 replication. Our results that these proteins had effects during infection with the ICP0-null but not WT virus indicated that their activities might be modulated by ICP0 or ICP0-induced processes to favor lytic infection. These regulatory pathways are likely additive to the known pathways, like the PML mediated pathway, which are targeted by ICP0 to promote lytic infection. SNX9 is a member of the sorting nexin family involved in intracellular trafficking. How it regulates HSV-1 replication is unknown. Like USP7, OTUD4 is a deubiquitinase. Our results showing that OTUD4 was induced by HSV-1 to enhance type I IFN production were consistent with a recent report that OTUD4 was induced by RNA viruses and targeted MAVS for deubiquitination thereby promoting signaling leading to type I interferon production [61]. The RIG-I like receptor RNA sensing pathway through MAVS is known to function during HSV-1 infection too [62]. Therefore it is possible that OTUD4 functions by a similar mechanism during HSV-1 infection although this hypothesis needs to be tested. VIM is an intermediate filament protein that participates in many biological processes including cytoskeletal assembly and immune response [63]. Although we found no evidence of its regulation of HSV-1 replication, it was reported to interact with NLRP3 and activate NLRP3 inflammasome in macrophages [64]. Consistent with its role in inflammation, VIM was reported to be upregulated in cerebralspinal fluid from children with enterovirus-associated meningoencephalitis [65]. Such functions in inflammation might not manifest in changes in virus production in cell culture and might require investigation in animal models.

In summary, the comprehensive analysis combining two proteomic approaches identified numerous potential interacting proteins and/or ubiqutination substrates of ICP0 in neuronal cells including previously known and novel ones. Several novel targets were validated. Functional studies of selective host targets revealed new regulators of viral replication and interferon response. This study helps understand the role of ICP0 in HSV-1 neuronal infection, and prompts further investigation of the targets identified in this study.

## Materials and Methods

### Cells and viruses

Vero, 293T, Neuro-2a and HFF were obtained from American Type Culture Collection and cultured as previously reported [45]. The method for culturing primary mouse TG neurons was decribed previously [38,44]. HSV-1 strains KOS, 7134, 7134R [66] and WT-BAC [47] were described previously. HSV-1 Flag-ICP0 virus was prepared according to a previously described protocol using bacterial artificial chromosome (BAC) technology [37,67]. Briefly, PCR-amplified zeocin-resistance (*Zeo*) cassette containing the DYKDDDDK (Flag)-coding sequence was inserted into one of the two copies of *ICP0* gene just downstream of the start codon. Then the PCR-amplified kanamycin-resistance (*Kan*) cassette containing the Flag-coding sequence and a homologous sequence was inserted into the other copy of *ICP0*. The *Kan* cassette was then removed by recombination via the homologous sequence resulting in markerless insertion of the Flag-coding sequence into the second *ICP0* gene copy. At the location of the first *ICP0* gene copy, the *Zeo* cassette was then replaced with the *Kan* cassette, which was subsequently removed as for the other *ICP0* copy. In each step correct colonies were selected by PCR and sequencing as well as restriction fragment length polymorphism following digestion with EcoRI and HindIII. BAC DNA was purified and transfected into Vero cells for Flag-ICP0 virus production. Primers used are listed in Table S3. HSV-1 infection of cells and plaque assays for viral titer measurements were performed as previously described [45].

### Plasmids

FLAG-HA-pcDNA3.1 (Addgene, #52535) was used to construct plasmids expressing HBB-BS, VIM, UL50, UBE2L3, C1QBP, Hist1H2BB, DDX17, HNRNPM, DBN1, PSMA6, P4HA2, MCM3, MCM5, RAP1A, OTUD4, SNX9, ACOT8, RPL10A, MYL12A, PPP1CB, CAVIN1 and PSMA3 with Flag-HA tags. HA-ubiquitin was constructed based on HA-pcDNA3.1 which was modified from FLAG-HA-pcDNA3.1. The firefly luciferase plasmid with IFN-β promoter (pIFN-β) and Renilla luciferase plasmid pRL-TK were kind gifts of Professor Chunfu Zheng at Fujian Medical University. ICP0 complete coding sequence was PCR-amplified from ICP0-WT plasmid (pRS-1) [68] and inserted into pcDNA3.1 without Flag or HA tags. Primers used for PCR were listed in Table S3.

### Transfection

All transfections were conducted using lipofectamine 3000 (Thermo Fisher) according to the manufacturer’s instructions. The siRNAs were synthesized by RiboBio. Their target sequences are: ACO8-si1, GACCCTAACCTTCACAAGA; ACOT8-si2, CCAAACAGATGTTCTGGGT; C1QBP-si1, CCACCTAATGGATTTCCTT; C1QBP-si2, CATTTGATGGTGAGGAGGA; OTUD4-si1, GCTTCTTCATGCTGAATAT; OTUD4-si2, CCAGCAGAACATATACCTT; SNX9-si1, GTGGGTTTATCCTACCTCT; SNX9-si2, CGGATCTATGATTACAACA; VIM-si1, CAGACAGGATGTTGACAAT; VIM-si2, CTTCTCAGCATCACGATGA

### Western blot analysis

Cells were scraped into sodium dodecyl sulphate (SDS) loading buffer (40% glycerol, 0.24 M Tris-HCl, pH 6.8, 8% SDS, 0.04% bromophenol blue, 5% β-mercaptoethanol) and heated at 95 °C for 5 min before being loaded onto SDS polyacrylamide gels for Western blot analysis. The following antibodies and dilutions were used: Flag antibody (MBL, M185-3L), 1:5000; ICP0 antibody (Santa Cruz, 11060), 1:2000; β-tubulin antibody (Sungene Biotech, KM9003), 1:5000; rabbit HA antibody (Sangon, D110004), 1:1000; β-actin antibody (Abclonal, Ac026), 1:5000; ICP4 antibody (Abcam, ab6514), 1:5000; gC antibody (Fitzgerald, 10-H25A), 1:1000; ICP27 antibody (Virusys, 1113), 1:5000; ACOT8 rabbit polyclonal antibody (Sangon, D221492), 1:500; C1QBP rabbit polyclonal antibody (Sangon, D152901), 1:1000; OTUD4 rabbit polyclonal antibody (Abclonal, A15229): 1:1000; Snx9 rabbit polyclonal antibody (Sangon, D154017), 1:1000; Vim rabbit polyclonal antibody (Sangon, D220268), 1:3000; HRP-conjugated goat anti-mouse (SouthernBiotech, 1030-05), 1:2000; goat anti-rabbit antibodies (SouthernBiotech, 4030-05), 1:2000.

### Sample preparation for ICP0 interactome profiling

2×10^7^ Neuro-2a cells in a T150 flask were infected with Flag-ICP0 or WT virus at an MOI of 20 and harvested at 6 hpi. Cells were first washed twice in ice-cooled PBS and then lysed in 1 mL of lysis buffer (50 mM HEPES-KOH [pH 7.4], 1% Triton X-100, 150 mM NaCl, 10% glycerol, and 2 mM EDTA plus one Complete EDTA-free protease inhibitor tablet [Roche] per 50 ml) for 1 hr. Supernatants were first incubated with 25 μL of mouse IgG for 1 hr at 4°C to remove the non-specific binding proteins and then incubated with 50 μL of Pierce Anti-DYKDDDDK Magnetic Agarose (ThermoFisher, A36797) for 2 hrs. The agarose was washed 4 times with lysis buffer for a total of 1 hr and then washed 4 times with PBS to remove Triton X-100. Proteins were eluted in 300 μL of elution buffer (0.1 M glycine, pH 2.8) at room temperature for 7 min and neutralized with 30 μL of Neutralization Buffer: (1 M Tris, pH 8.5). Experiments in HFF and 293T cells were conducted in the same way except that an MOI of 10 was used for infection. Immunoprecipitated proteins were reduced with 10 mM dithiothreitol (DTT) for 45 min at 30°C and alkylated with 30 mM iodoacetamide (IAA) for 30 min at room temperature in the dark. Protein were then cleaned up by acetone precipitation and resuspended in 50 mM ammonium bicarbonate. Protein was digested by trypsin (Promega) at a ratio of 1:50 (trypsin:protein) at 37°C overnight. Tryptic peptides were acidified by trifluoroacetic acid (TFA) and was desalted using a HLB microplate (Waters). The microplate was conditioned with 200 μL acetonitrile (ACN), followed by 200 μL 60% ACN, then by 200 μL 0.1% TFA twice. The sample was loaded to the microplate and then washed three times with 200 μL 0.1% TFA. Desalted peptides were eluted with 60% ACN and lyophilized in a vacuum lyophilizer (LABCONCO) before LC-MS/MS analysis.

### Sample preparation for quantitative ubiquitinome investigation

The following three groups were compared to determine the effects of ICP0 on ubiquitinome: Group A: pcDNA3.1 transfection + ICP0 null virus infection; Group B: ICP0 plasmid transfection + ICP0 null virus infection and group; Group C: pcDNA3.1 transfection + ICP0-rescued virus infection. Neuro-2a cells in a 100-mm dish were transfected with ICP0 or pcDNA3.1 plasmid for 24 h before infection with 7134 virus (ICP0-null) or 7134R virus (ICP0-rescued) at an MOI of 20. After the one-hour incubation for virus absorption, fresh medium was added with 10 μM MG-132 to inhibit proteasome-dependent degradation. The cells were harvested at 6 hpi in urea lysis buffer (20 mM Hepes, pH 8.0, 9 M urea). After sonication and centrifugation, the supernatants were kept.

Proteins (10 mg) were reduced with 10 mM dithiothreitol (DTT) for 45 min at 30°C and subsequently alkylated with 30 mM iodoacetamide (IAA) for 30 min at room temperature in the dark. Lysis buffer was then replaced by 50 mM triethylammonium bicarbonate (TEAB) using Zeba Spin Desalting Columns. Protein was digested by TPCK-treated trypsin (Sigma) at ratio of 1:40 (trypsin:protein) at 37°C overnight. Tryptic peptides were acidified by trifluoroacetic acid (TFA) and was desalted using a 500 mg C18 Sep-Pak SPE cartridge (Waters). C18 cartridges were conditioned with 4 mL acetonitrile (ACN), followed by 4 mL 50% ACN+0.1% TFA, then by 4 mL 0.1% TFA. The sample was loaded to the C18 cartridge and then washed three times with 4 mL 0.1% TFA. Desalted peptides were eluted with 50% ACN+0.1% FA and lyophilized in a vacuum lyophilizer (LABCONCO).

### Enrichment of K-ε-GG peptides

Di-Gly modified peptides derived from ubiquitinated proteins were enriched by PTMScan ubiquitin remnant K-ε-GG motif kit (CST) according to the manufacturer’s instruction. Briefly, antibody-loaded beads were washed three times with 1 mL pre-cooled 100 mM sodium borate (pH 9.0). Antibody was then cross-linked by 20 mM dimethyl pimelimidate for 30 min. Cross-linked antibody beads (50 μg) were mixed with tryptic peptides in cold 1× IAP buffer for 2 h at 4°C with end-over-end rotation, followed by three time ice-cold IAP buffer washes and an ice-cold water wash. DiGly-modified peptides were eluted by 50 μl 0.15% TFA twice and dried completely by vacuum centrifugation.

### TMT isobaric labeling

DiGly-modified peptides enriched from each sample were dissolved in 20 μL 100 mM TEAB and labeled with 0.8 mg 10plex TMT Label Reagent (ThermoFisher) in 50 μL ACN. The reaction was proceeded at room temperature for 1 hour and stopped by 2 μL 5% hydroxylamine. Supernatant was desalted by homemade C18 stage tip and dried by vacuum centrifugation.

### StageTip-based strong cation-exchange (SCX) fractionation of K-ε-GG peptides

Home-packed strong cation-exchange chromatography StageTip with Empore Cation-SR Extraction material (3M) was sequentially conditioned by 100 μL ACN, 80% ACN+0.1% TFA and 5% ACN+0.1% TFA. The combined TMT-labeled K-ε-GG peptides were resuspended in 80 μL 5% ACN+0.1% TFA and loaded to SCX Stagetip. The flow-through was labeled as SCX fraction 1. Peptide were then sequentially eluted with 100 μL 5% ACN+0.1% TFA supplemented with increasing concentration of ammonium acetate (40, 100, 200, 350, 500 mM). The corresponding eluates were designated as SCX fractions 2, 3, 4, 5 and 6, respectively. All peptide fractions were dried by vacuum centrifugation.

### LC-MS/MS analysis

All dried peptides were resuspended in 2% ACN and 0.1% FA and separated by nanoLC-MS/MS using an UltiMate 3000 RSLCnano system (Thermo Scientific) at the flow rate of 400 nL/min. Solvent A is 2% ACN, 0.1% FA and solvent B is 98% ACN, 0.1% FA. Gradient elution was performed at 50°C using linear gradients of 120 min: 1-4 min with 3% (v/v) of s B, 4-6 min from 3% to 5% (v/v) B, 6-70 min from 5% to 15% (v/v) B, 70-90 min from 15% to 30% (v/v) B, 90-100 min from 30% to 80% (v/v) B, 100-110 min with 80% (v/v) B, 110-120 min with 3% (v/v) A. The eluted peptides were analyzed by Q Exactive HF-X (Thermo Scientific) acquiring MS spectra at the resolution of 120,000 FWHM with a mass range of 300–1500 m/z and an AGC target of 3E6. The top 20 precursors were then fragmented by HCD with collision energy approximately 32% NCE and MS2 spectra were acquired at the resolution of 45,000 FWHM.

### Mass spectrometry data analyses

All MS raw data were loaded into MaxQuant (version 1.6.2.10) and searched against human or Mouse UniProtKB database (November 2019) supplemented with HSV1 sequences, with the automatic reverse database on target-decoy search mode. MS2 based isobaric labeling using 10plex TMT tags were configured only for the ubiquitinome analysis. Variable modifications included oxidation (M) (+15.99491 Da) and acetyl (protein N-term) (+42.01056 Da) were specified for ubiquitinome, proteome and ICP0-interactome analyses, while GlyGly(K)_10plex_TMT (+343.20586 Da) was specified only for the ubiquitinome analysis. Carbamidomethyl (C) (+57.02146 Da) was set as the fixed modification. Trypsin was set as digestion mode with 2 maximum missed cleavage sites. We used 20 ppm in the first search ion tolerance and 4.5 ppm in the main search ion tolerance. MaxQuant default setting was used for all other parameters. Peptide and protein identification were both filtered by FDR < 1%.

### Statistical and functional enrichment analyses

For the interactome data sets from 293T and Neuro-2a cells, potential ICP0 interacting proteins were selected by either fold change ≥ 5, or fold changes ≥ 1.2 with p < 0.05 (t test) between Flag-ICP0 and WT comparison. For HFF cells, which identified a larger number of proteins, we used a more stringent criterion of fold changes > 5 and p < 0.1 (t test). Proteins with <2 unique peptides were discarded. The ubiquitinome data were normalized by protein levels determined by peptides without di-glycine modification were used to normalize. After normalization, modified sites with > 1.25 higher di-glycine levels in both B/A and C/A comparisons, or with increasing di-glycine levels with FDR<0.05 (t tests) were considered as potential ICP0 target sites. To identify host pathways modulated by HSV-1 ICP0, potential ubiquitination targets of ICP0 were then submitted for functional annotation enrichment analysis in KOBAS 3 [69] (http://kobas.cbi.pku.edu.cn) against Reactome database [70]. Fisher’s exact test and Benjamini and Hochberg’s FDR <0.01 method were chosen for significant enriched terms. Enriched term with >1000 genes in database or <10 mapped genes were discarded from analyses. Similar or redundant terms were manually removed to keep the most significantly enriched ones. Heat map was painted using the heatmap3 package in R software. All potential ICP0 interacting proteins were used to generate interaction network based on String database (https://string-db.org) [71] while all the potential ubiquitination targets were submitted to search conserved domains by InterPro of String database [72].

### Co-immunoprecipitation (Co-IP) for validating interactions

Five hours after 5 x 10^6^ Neuro-2a cells were plated in 100-mm dishes, the cells were transfected with the ICP0 expressing plasmids together with one of the plasmids expressing a potential interacting protein with a Flag tag. Cells were lysed in 1 mL lysis buffer (see above) at 30 hpt. The Flag-tagged proteins were immunoprecipitated as described for the ICP0 interactome analysis except that 40 μL anti-DYKDDDDK Magnetic Agarose was used and 80 μL 1 x SDS loading buffer was added to elute proteins. Western blotting was conducted to detect the proteins.

### Validation of ubiquitination

About 9 x 10^6^ Neuro-2a cells in 100-mm dishes were co-transfected with the following plasmids: 5 μg pOTUD4 or pVIM (Flag-HA tag), 2μg HA-Ubiquitin and 3 μg ICP0 or ICP0-RFm as control. Cells were lysed in lysis buffer at 48 hpt. IP was performed as described above using the anti-Flag agarose.

### Quantitative reverse-transcription-PCR

To quantify mRNA levels in cells, total RNA was purified using a RNA Isolation Kit (catalogue #. RC112, Vazyme Biotech) according to the manufacturer’s instructions. Reverse transcription and PCR were conducted using a HiScript II Q Select RT SuperMix and ChamQ Universal SYBR qPCR Kit (catalogue #. R233-01 and Q711-02/03, Vazyme Biotech). Data was normalized to GAPDH mRNA levels. The following primers for mouse transcripts were used. GAPDH-F: GAAGGTCGGTGTGAACGGATT; GAPDH-R: GCCTTGACTGTGCCGTTGAA; OTUD4-F: CCTCCATCTCAGGTGTCTGAAGGTCA; OTUD4-R: GGTTAGGCCCAAAAGACTGTTGTGG; IFN-α-F: CTCATTCTGCAATGACCTCCACC, IFN-α-R: ACTTCTGCTCTGACCACCTCCC

### Luciferase assays

Neuro-2a cells were co-transfected with 400 ng/ml of the indicated plasmid or 80 nM of the indicated siRNA, 80 ng/ml of a pIFN-β and 40 ng/ml of pRL-TK for 48 h, then infected with 7134 or KOS virus at an MOI of 2 for the indicated times. The cells were then lysed and promoter activities were measured by a dual luciferase kit according to the manufacturer’s protocol (Yeasen, 11402ES60).

## Data availability

Raw LC-MS/MS data of both interactome and ubiquitinome analyses were stored in the public proteome data resources (iProX ID: IPX0002267000, ProteomeXchange ID: PXD019782), and can be accessed via the following address: https://www.iprox.cn/page/PSV023.html;?url=1639049337236KEof (password:irow).

## Acknowledgments

We thank the Core Facility of Zhejiang University School of Medicine for expertise and instrument availability. This work was supported by the National Key R & D program of China (2017YFC1200204) to D.P. and Z.S., National Natural Science Foundation of China (81671993) to D.P and (U20A20343) to Z.S., Natural Science Foundation of Zhejiang Province (LR18H190001) to D.P.

## Competing Interests

The authors have declared that no competing interests exist.

---

**Fig. S1.**
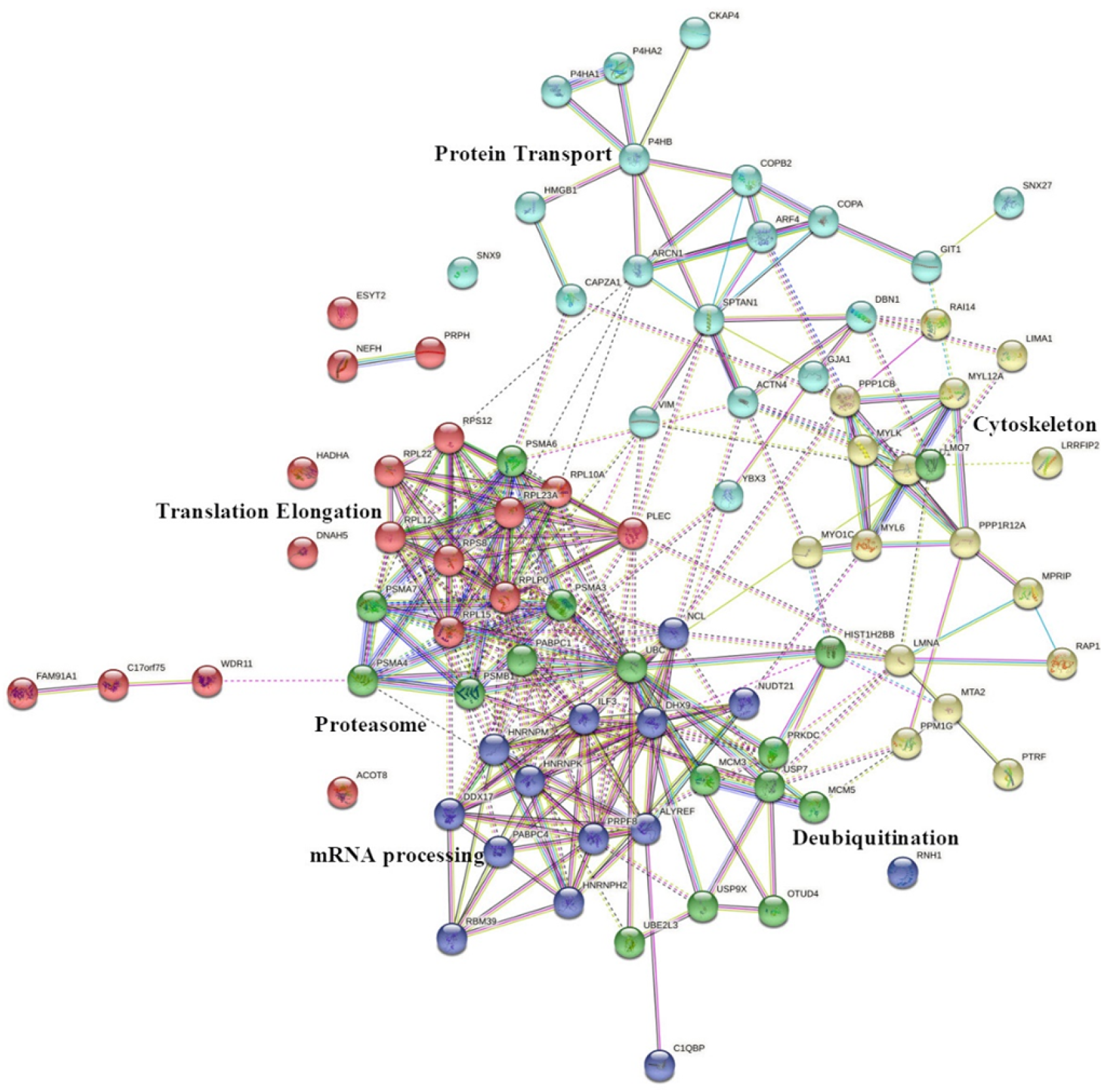
ICP0-interacting network. The functional clusters were generated by the K-mean approach based on the candidate ICP0-interacting proteins identified by the interactome analysis in all three cell lines.

**Fig. S2.**
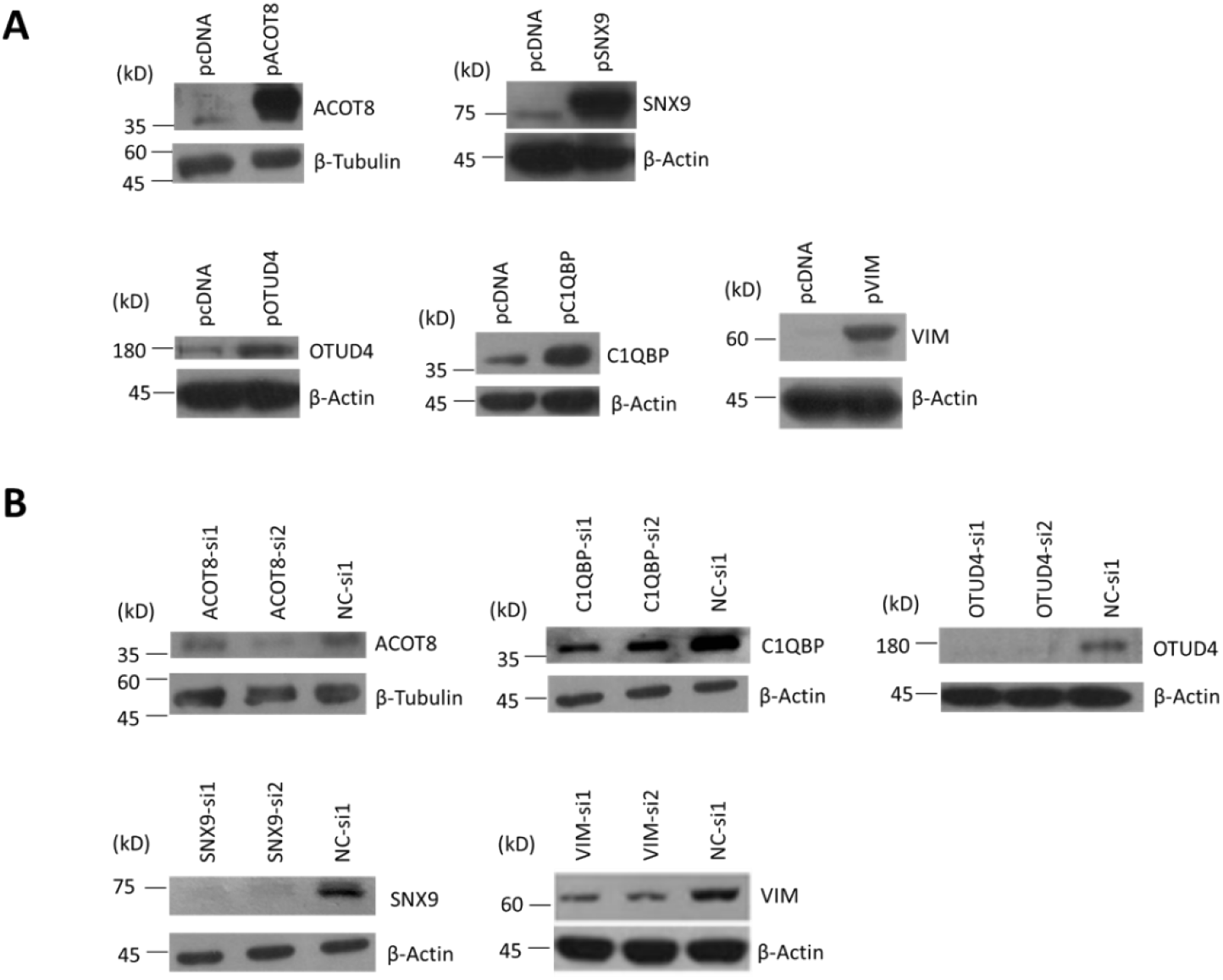
Validation of overexpression and knockdown of ICP0-interacting proteins in Neuro-2a cells. (A) Neuro-2a cells were transfected with 400 ng/ml of the indicated plasmids for 48 h before being harvested for western blot analysis for the indicated proteins. (B) Neuro-2a cells were transfected with 120 nM of the indicated siRNA for 48 h before being harvested for western blot analysis. NC, negative control.

**Fig. S3.**
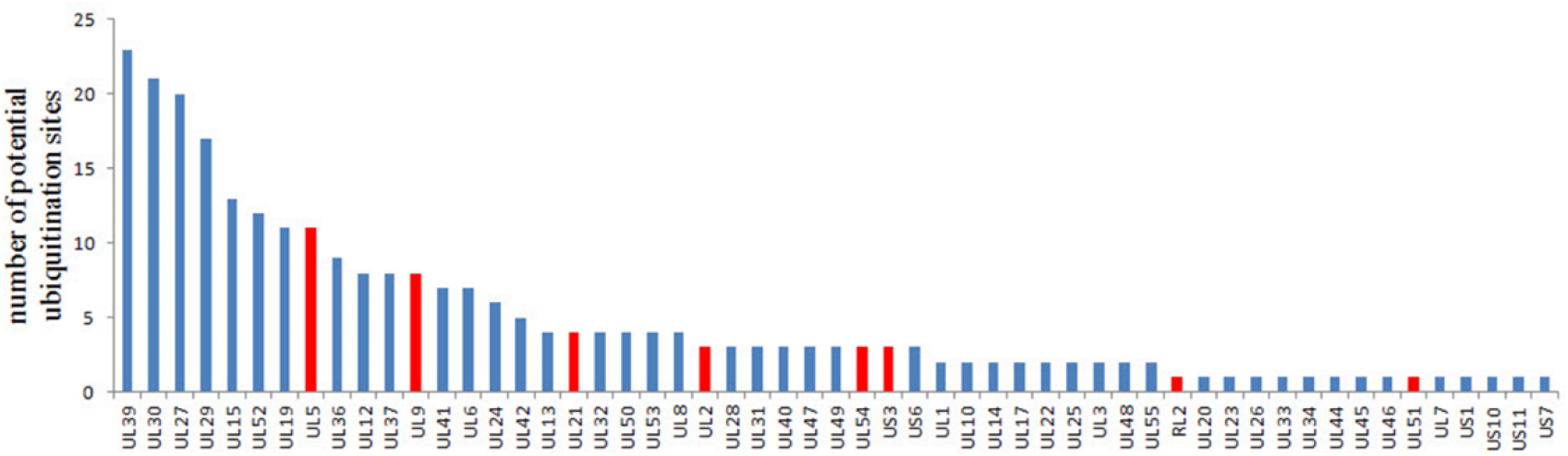
Potentially ubiquitinated viral proteins as detected by mass spectrometry. Red color indicates proteins potentially targeted by ICP0 because their ubiquitination levels were increased by ICP0 presence.

**Table S1. Processed data for all MS data used in this study. This table is provided as an attached Excel file.**

**Table S2.** Potential ICP0 substrates identified by ubiquitinome analysis classified by superfamilies according to conserved domains.

| Term description | Matching proteins |
| --- | --- |
| P-loop containing nucleoside triphosphate hydrolase | DDX3X, MPP2, DLG4, SMC6, GTPBP4, PSMC1, WRNIP1, PSMC6, HELLS, EIF4A3, NVL, ORC4, VCP, GNL2, ABCF2, DDX54, WRN, SRPR, ASCC3, HNRNPUL1, DHX9, EEF2, KIF2C, ATL2, DDX10, DDX1, ERCC6, UPF1, SMC5, CHD8, RNF213, DDX24, GNL3L, KIF3A, LSG1, CHD2, BLM, KATNA1, KIF20A |
| Zinc finger, RING/FYVE/P HD-type | UHRF1, RUFY1, UHRF2, HGS, RNF34, LTN1, BAZ1A, USP5, TRAFD1, RNF149, WHSC1, ZMYND11, PML, RNF213, TTC3, USP33, RNF168, BAZ2A |
| Nucleotide-binding alpha-beta plait domain superfamily | ELAVL3, RBM28, G3BP1, NCBP3, NIFK, RBM17, RALY, UPF3B, U2SURP, PABPC4, TARDBP, RBM27, HNRNPC, CELF4, SRSF6, PTBP1, PARP10, HNRNPF |
| Armadillo-type fold | IFRD1, UTP20, CTNNB1, CEBPZ, HUWE1, PSMD1, PSMD5, SDAD1, PI4KA, LTN1, THADA, MYBBP1A, BZW1, EIF4G3, CLTC, XPO1, RALGAPA1, PUM2 |
| RNA-binding domain | ELAVL3, RBM28, G3BP1, NIFK, RBM17, RALY, UPF3B, U2SURP, PABPC4, TARDBP, RBM27, HNRNPC, CELF4, SRSF6, PTBP1, HNRNPF |
| DEAD/DEAH box helicase domain | DDX3X, EIF4A3, DDX54, WRN, ASCC3, DHX9, DDX10, DDX1, DDX24, BLM |
| Ubiquitin interacting motif | ATXN3, HGS, STAM, UIMC1 |
| Ubiquitin specific protease domain | USP15, USP43, USP11, USP5, USP33, USP7 |

**Table S3.**
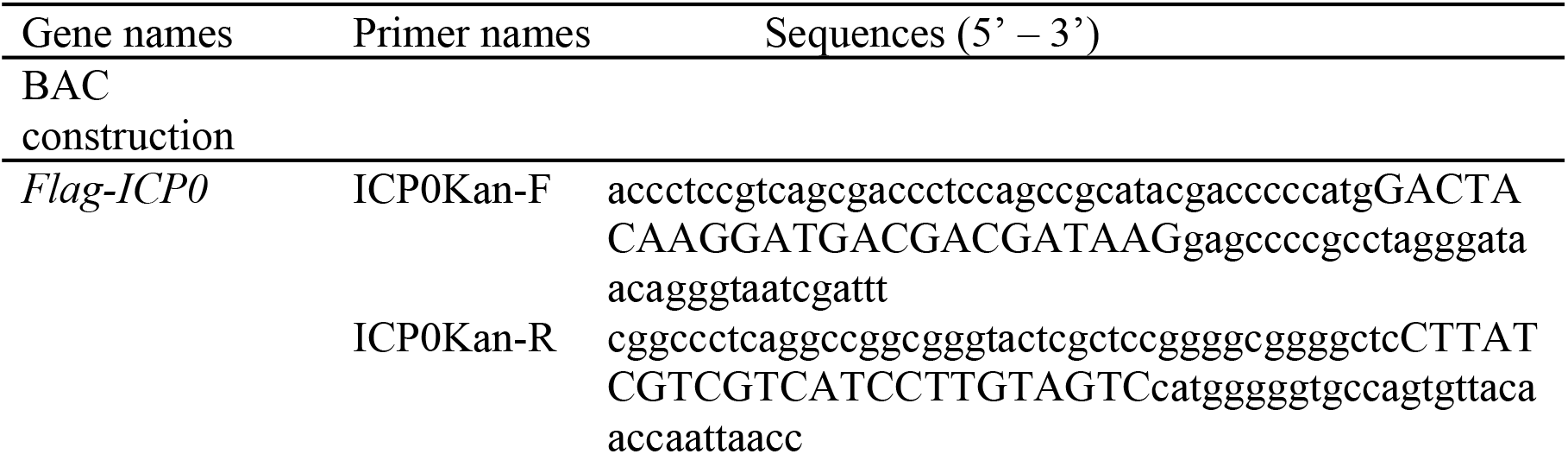
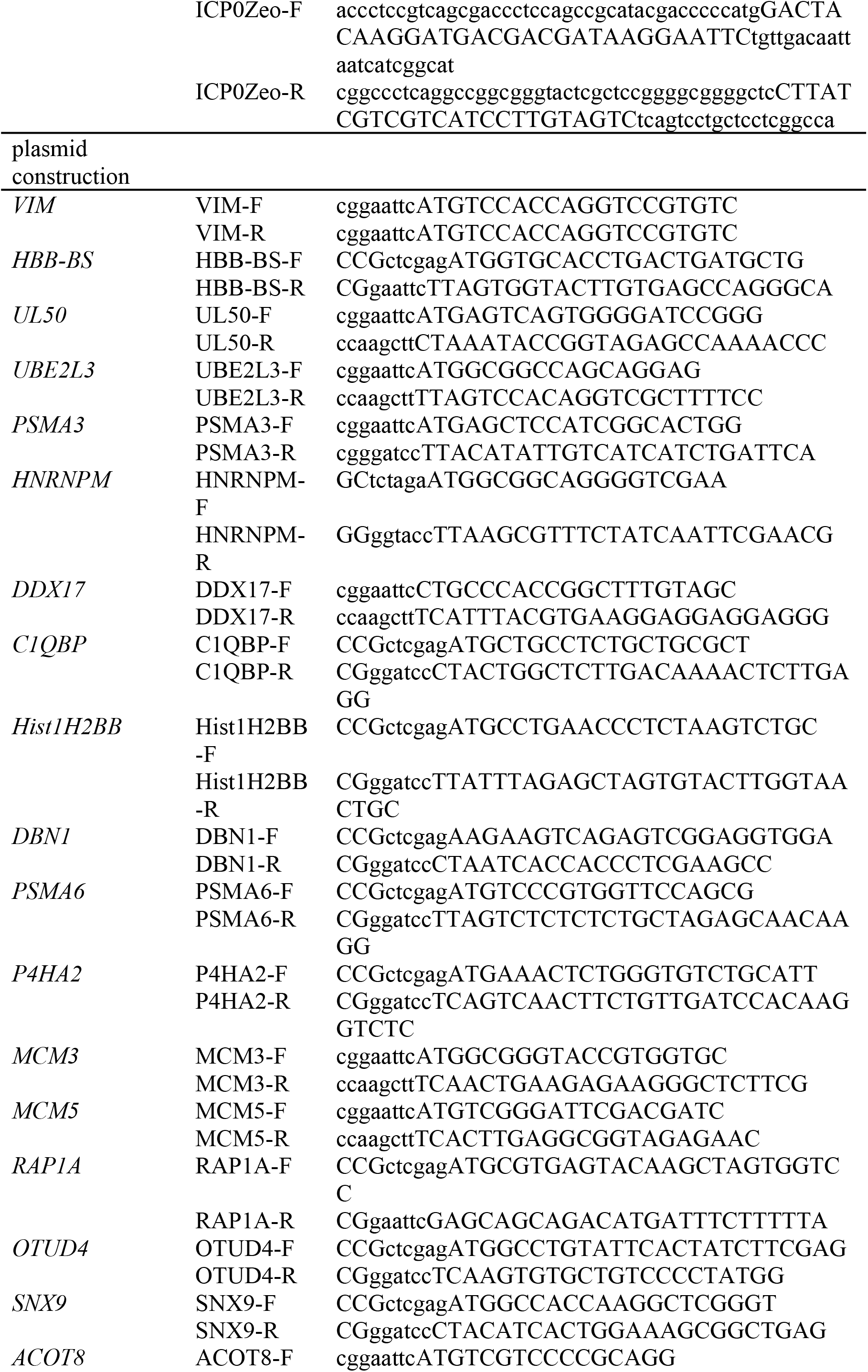
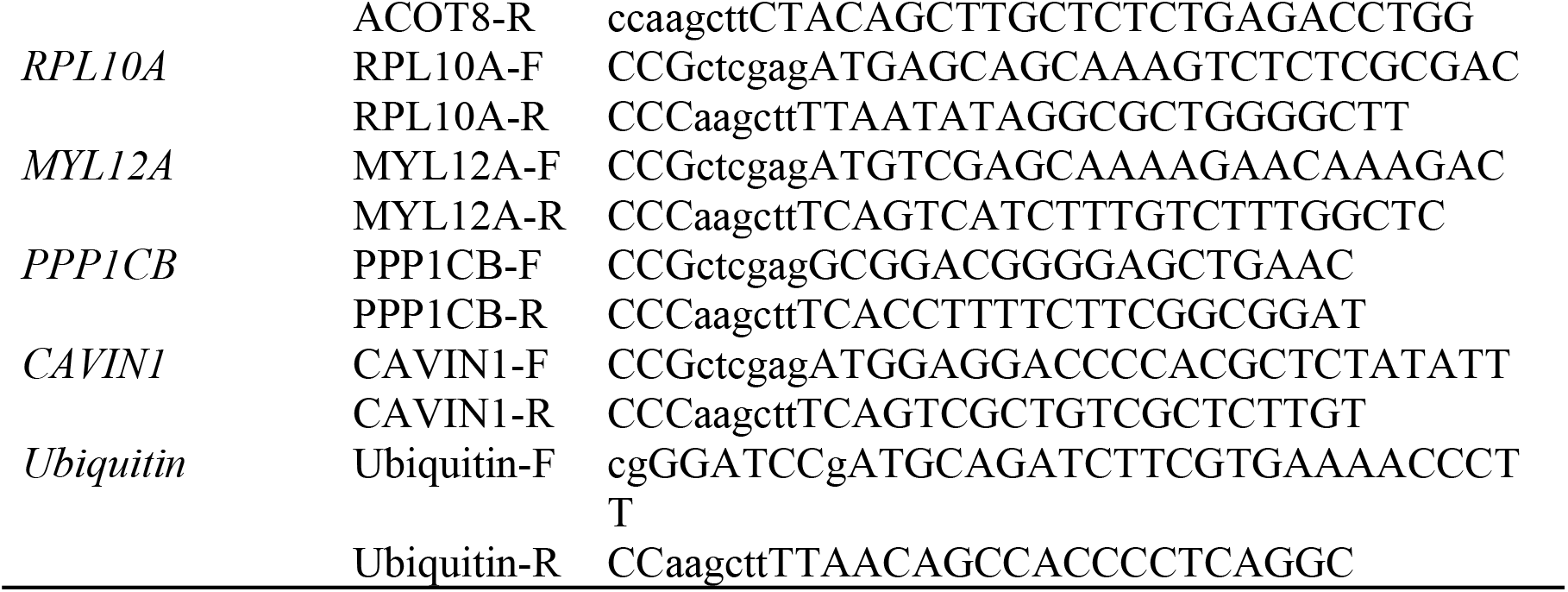
Primers used in this study.

## Notes

### Competing Interest Statement

The authors have declared no competing interest.

